# Mediator subunit MED16 cooperates with UBP1-TFCP2 to control the promoter context-dependent transcriptional activity

**DOI:** 10.1101/2025.08.12.669905

**Authors:** Yuan-Ming Zheng, Xia-Ying Zhao, Ming Yang, Xin-Yi Yang, Huan-Zhang Zhu, Xiao-Fei Yu, Qiang Zhou, Gang Wang

## Abstract

The Mediator complex acts as a master transcriptional co-regulator, yet the mechanisms by which its individual subunits mediate gene-specific transcription remain incompletely defined. Here, we report that the Mediator subunit MED16 physically associates with the UBP1-TFCP2 heterodimer, which controls the context-dependent transcriptional activation or repression. Biochemical purification combined with mass spectrometry validated MED16 as a bona fide interactor of UBP1-TFCP2. Integrative transcriptomic analyses demonstrated that UBP1 recruits the Mediator complex through MED16 to activate a panel of silenced genes associated with lung homeostasis, angiogenesis and cell proliferation. By contrast, MED16-UBP1 co-occupancy at the HIV-1 transcription start site (TSS) represses viral transcription and sustains HIV latency by compromising pre-initiation complex assembly. Domain-mapping experiments revealed that MED16 must integrate into the intact Mediator complex to exert either activating or repressive function. Genomic analysis uncovered a positional regulatory logic: UBP1-TFCP2 binding motifs adjacent to the TSS promote transcription, while those overlapping the TSS elicit repression. Collectively, our work identifies an MED16-UBP1 regulatory module with dual transcriptional activity and highlights this signaling axis as a potential target for HIV-1 interventions.

## Introduction

Transcription by RNA polymerase II (Pol II) requires the assembly of a pre-initiation complex (PIC) at gene promoters. Among the co-activators that facilitate this process, the multi-subunit Mediator complex plays a central role (1–8). The Mediator complex is composed of ∼30 subunits in mammals and structurally divided into four modules: Head, Middle, Tail, and Kinase. The scaffold subunit MED14 joins the Tail, Middle, and Head modules together to form the “core Mediator”. The tail module is composed of MED15, MED16, MED23, MED24, MED25, MED27, MED28, MED29, and MED30; and the main function of the tail module is to provide an interactional platform for transcription factors to recruit Mediator complex to gene promoters and enhancers for further regulation of gene expression (8–10). Among all tail subunits, MED16, MED23, MED24, and MED25 form a submodule that accounts for ∼70% molecular weight of the tail module and creates a large platform for interactions of transcription factors (11). Previous studies have found MED23 interacts with multiple transcriptional factors and controls diverse gene expression programs at multiple stages of transcription processes (12–15). It prompts us to question whether other subunit in this submodule play similar or distinctive roles in transcriptional regulation, especially MED16 that is structurally conserved from yeast to mammals and acts as a critical regulator in yeast and plants. Sin4 (yeast MED16) plays both positive and negative roles in the transcriptional regulation of many genes in yeast, and loss of Sin4 alters chromatin accessibility globally (16). In plant cells, MED16 controls endoreduplication and cell growth by repressing the expression of APC/C activator genes *CCS52A1/A2* (17–21). MED16 also controls plant cold response, chemical and wound response, and iron homeostasis at transcriptional levels (17–21). However, the functions of MED16 in mammalian cells are poorly understood. Among a few reports, MED16 was reported as a key factor in NRF2-activted gene expression via its direct interaction with NRF2. Knockout of MED16 reduces NRF2-dependent anti-oxidative gene expression (21, 22). Additionally, MED16 was down-regulated in papillary thyroid cancer, leading to increased TGF-β signaling and radioiodine resistance (23). Though the exact molecular mechanisms by which MED16 regulates gene expression remain to be further elucidated, and its roles in animal development and diseases are barely investigated.

UBP1 (as known as LBP-1) and TFCP2 (as known as LSF) belong to a subfamily of the TFCP2/Grainyhead transcription factors family. The amino acid sequences of TFCP2 and UBP1 share 88% identity and they can form both homo-and heterodimers (24–26). UBP1 and TFCP2 bind to the consensus sequence CNRG-N6-CNR(G/C) (27). They function as a transcription activator or repressor to regulate a number of viral and cellular genes, including mouse α*-Globin*, *MMP-9*, *CYP11A1*, *IL-4*, SV-40, and HIV-1 (28–32). UBP1 and TFCP2 are involved in various biological processes, including embryonic development, angiogenesis, tumorigenesis, and HIV-1 latency (28, 29, 31–38). There is a canonical UBP1 binding motif within the HIV-1 core promoter which overlaps with the transcription start site (TSS). UBP1 has been implicated in HIV-1 transcriptional repression through several reported mechanisms, including elongation restriction and recruitment of YY1/HDAC1 (32–34). However, how UBP1 and TFCP2 regulate cellular gene expression, and what are the genome-wide target genes of UBP1 and TFCP2 still remain unclear.

In this study we identified an interaction between MED16 and UBP1-TFCP2 dimer utilizing gel-filtration chromatography and immunoprecipitation-mass spectrometry (IP-MS). Domain mapping revealed that MED16 uses its N-terminal WDR domain to anchor within the Mediator complex and its C-terminal αβ-domain to bind UBP1. Functionally, MED16 must act through the Mediator complex to support UBP1-mediated transcriptional activation, including the activation of pulmonary surface molecule genes. Conversely, MED16 cooperates with UBP1 within the Mediator complex to inhibit HIV-1 transcription at the viral transcription start site. At the genomic scale, UBP1 activates gene expression when the UBP1-TFCP2 binding motif is located near TSS, but inhibits gene expression by obstructing normal PIC formation when the UBP1-TFCP2 binding motif overlaps with TSS. Therefore, the position of the UBP1 binding site relative to the TSS determines whether UBP1-MED16 activates or represses transcription. These findings provide mechanistic insights into UBP1-mediated HIV-1 repression and suggest a potential strategy for HIV-1 therapy.

## RESULT

### A fraction of MED16 biochemically dissociates from the core Mediator complex

Multiple Mediator subunits consist of large intrinsically disordered regions (IDRs) and lacks predictable functional motifs (39, 40). The Mediator complex forms liquid-liquid phase separation condensates at super-enhancers. These IDR-mediated condensates can be disrupted by 1,6-hexanediol (1,6-HD) (41, 42), an aliphatic alcohol, via inhibition of weak hydrophobic protein-protein interactions (43–47).

To determine whether individual Mediator subunits exhibit differential sensitivity to hydrophobic perturbation, we immunoprecipitated endogenous Mediator complexes from 293T cell lysates using antibodies against CDK8, MED1, and MED12 respectively, followed by washing with buffers containing increasing concentrations of 1,6-HD. Most of the detected subunits show modest decreases, whereas MED16 exhibits the most pronounced reduction in response to increasing 1,6-HD treatment (Fig 1A). To validate this observation, we performed parallel experiments using HeLa nuclear extracts (NEs) with 2,5-hexanediol (2,5-HD) as a negative control, as 2,5-HD does not disrupt weak hydrophobic interactions like 1,6-HD does. Consistently, MED16 dissociation was specifically induced by 1,6-HD but not by 2,5-HD in HeLa nuclear extract (Fig 1B).

**Figure 1.**
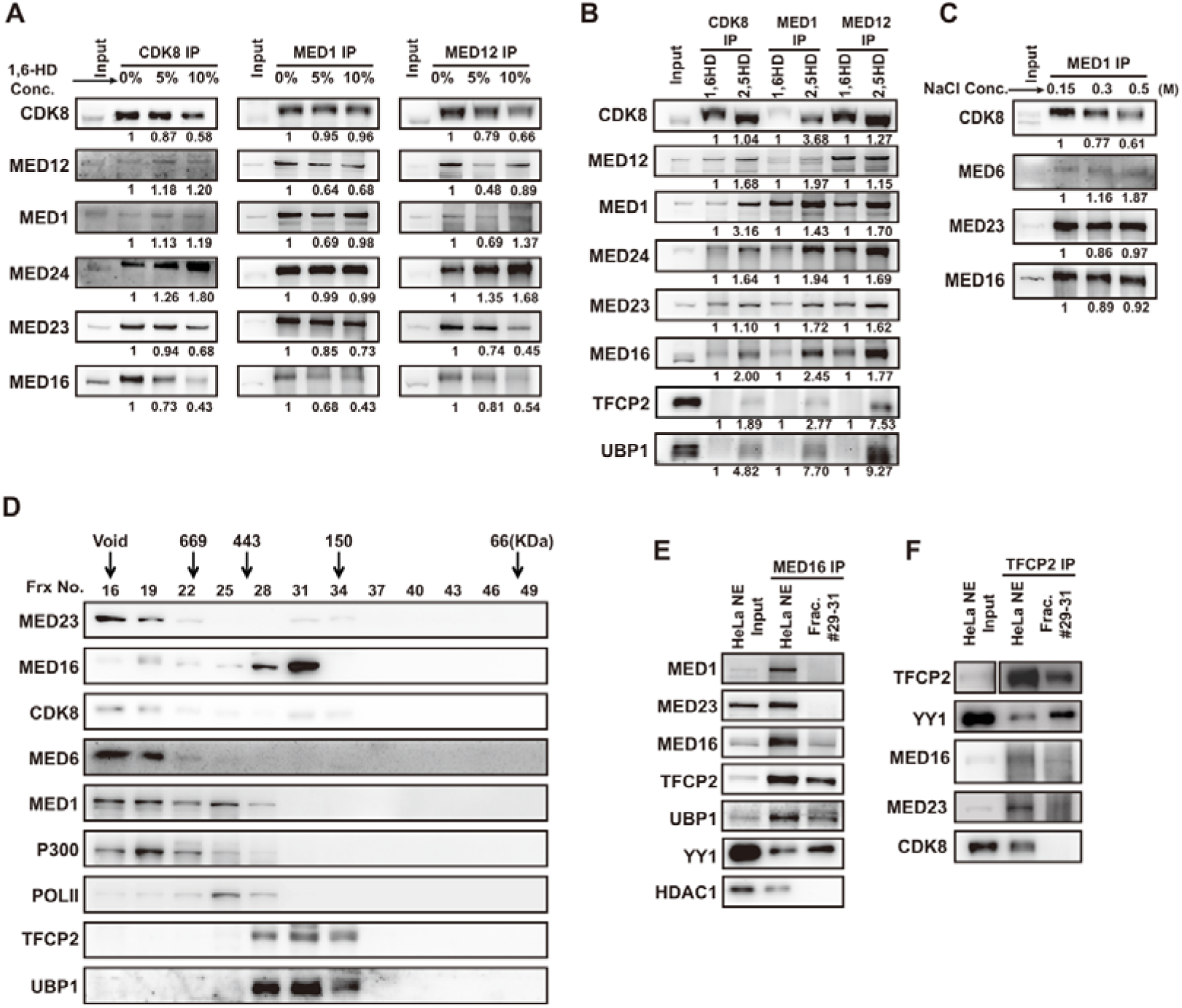
Gel-filtration and 1,6-hexanediol treatment resolve a MED16-containing fraction that interacts with UBP1-TFCP2. (A) Co-IP experiment with antibodies against endogenous CDK8, MED1, and MED12 in 293T whole cell lysis. The resultant immunoprecipitates were washed by washing buffers containing 0%, 5%, and 10% of 1,6-hexanediol respectively. Input: 0.5% of the total lysate was loaded. (B) Co-IP experiment with antibodies against endogenous CDK8, MED1, and MED12 in HeLa nuclear extract. The resultant immunoprecipitates were washed by washing buffers containing 10% of 1,6-hexanediol or 10% 2,5-hexanediol. Input: 0.2% of the total HeLa nuclear extract was loaded. (C) Co-IP experiment with antibodies against endogenous MED1 in 293T whole cell lysis. The resultant immunoprecipitated was washed by washing buffers containing 150 mM, 300 mM, and 500 mM of NaCl respectively. Input: 0.2% of the total HeLa nuclear extract was loaded. (D) Gel filtration chromatography of HeLa nuclear extract. 500µl HeLa nuclear extract was applied to Superose 6 column then was run in Buffer D. Column fractions of 500µl were collected. The Void (void volumn) of Superose 6 is based on the volume of effluent required for the elution of blue dextran (molecular mass of ∼2000 kDa). The molecular weight of corresponding fractions was detected by protein standards. (E) Immunoblots of anti-MED16 immunoprecipitation in HeLa nuclear extract and gel filtration factions No.29 to No.31. The immunoprecipitated proteins were detected with indicated antibodies by western blotting. HeLa NE input: 0.2% of the total HeLa nuclear extract was loaded. (F) Immunoblots of anti-TFCP2 immunoprecipitation in HeLa nuclear extract and gel filtration factions No.29 to No.31. The immunoprecipitated proteins were detected with indicated antibodies by western blotting. HeLa NE input: 0.2% of the total HeLa nuclear extract was loaded.

We next tested whether the MED16-Mediator interaction is mediated by ionic contacts by performing co-IP under increasing salt concentrations. Immunoprecipitated Mediator complexes were washed with buffers containing 150 mM, 300 mM, and 500 mM NaCl respectively. MED16 binding to the core complex remained unaltered under high-salt conditions (Fig 1C), indicating that MED16 integrates into the Mediator complex primarily through hydrophobic rather than electrostatic interactions. In parallel, we analyzed the elution profile of MED16 by gel-filtration chromatography of HeLa NE. Most Mediator subunits co-eluted at ∼2,000 kDa, corresponding to the intact holo-Mediator complex (Figure 1D). Notably, a substantial fraction of MED16 was also detected in lower molecular weight fractions (between 443 kDa and 150 kDa), well separated from the peak of core Mediator subunits (Figure 1D). Together, the 1,6-HD sensitivity and high-salt resistance indicate that MED16 associates with the Mediator complex primarily through hydrophobic interfaces. In addition, gel-filtration fractionation reveals that a portion of MED16 can be separated from the core Mediator complex in vitro.

### UBP1 and TFCP2 associate with Mediator complex via MED16 subunit

MED16 has a molecular weight of approximately 95 kDa, but gel-filtration fractionation showed that the dissociated MED16 in fractions corresponds to 443-150 kDa (Fig 1D). This suggested that it may associate with other proteins. To identify these potential partners, we performed immunoprecipitation-coupled mass spectrometry (IP-MS) using antibodies against MED16, MED1, MED12, and CDK8 respectively. Two dimeric transcription factors, UBP1 and TFCP2, were identified as preferential MED16-binding proteins (Fig S1). Their co-elution with MED16 in the same gel-filtration fractions was confirmed by immunoblotting (Fig 1D). This suggests that UBP1 and TFCP2 may interact with the MED16 subunit (Fig S1).

To validate the association between UBP1/TFCP2 and MED16, an antibody specific to MED16 was used to capture the MED16 associated proteins in HeLa NE or in the 443-150 kDa fractions. In HeLa NE, MED1, MED23, TFCP2 and UBP1 were co-immunoprecipitated with anti-MED16 antibody, but only TFCP2 and UBP1 were co-immunoprecipitated with MED16 in 443-150 kDa fractions (Fig 1E). Reciprocally, anti-TFCP2 antibody is also able to pull down MED16 in either HeLa NE or 443-150 kDa fractions (Fig 1F). YY1 and HDAC1 are known to complex with TFCP2 and UBP1 to suppress HIV-1 transcription (34, 48, 49). YY1 was pulled down in both samples, whereas only a small amount of HDAC1 was pulled down from HeLa NE but not from the 443-150 kDa fractions (Fig 1E).

The above experiments established that UBP1 and TFCP2 interact with the biochemically resolved MED16. We next asked whether these transcription factors can also associate with the intact Mediator complex. To test this, we re-probed the IP samples from Figure 1B with antibodies against UBP1 and TFCP2. Both proteins were co-immunoprecipitated with the Mediator complex under 2,5-HD conditions, and this association was lost upon 1,6-HD treatment, which disrupts the MED16-Mediator interface (Fig 1B). These results indicate that UBP1 and TFCP2 can associate with the intact Mediator complex, likely through MED16.

To confirm that UBP1-TFCP2 associate with the Mediator complex specifically through MED16, we next mapped the functional domains responsible for MED16-UBP1/TFCP2 interactions. MED16 contains an N-terminal WD40-repeat (WDR) domain and a C-terminal αβ-domain. Serial truncation mutagenesis revealed that the N-terminal WDR domain is essential for MED16 integration into the core Mediator complex, whereas the C-terminal αβ-domain is strictly required for binding to UBP1-TFCP2 (Fig 2A, 2B).

**Figure 2.**
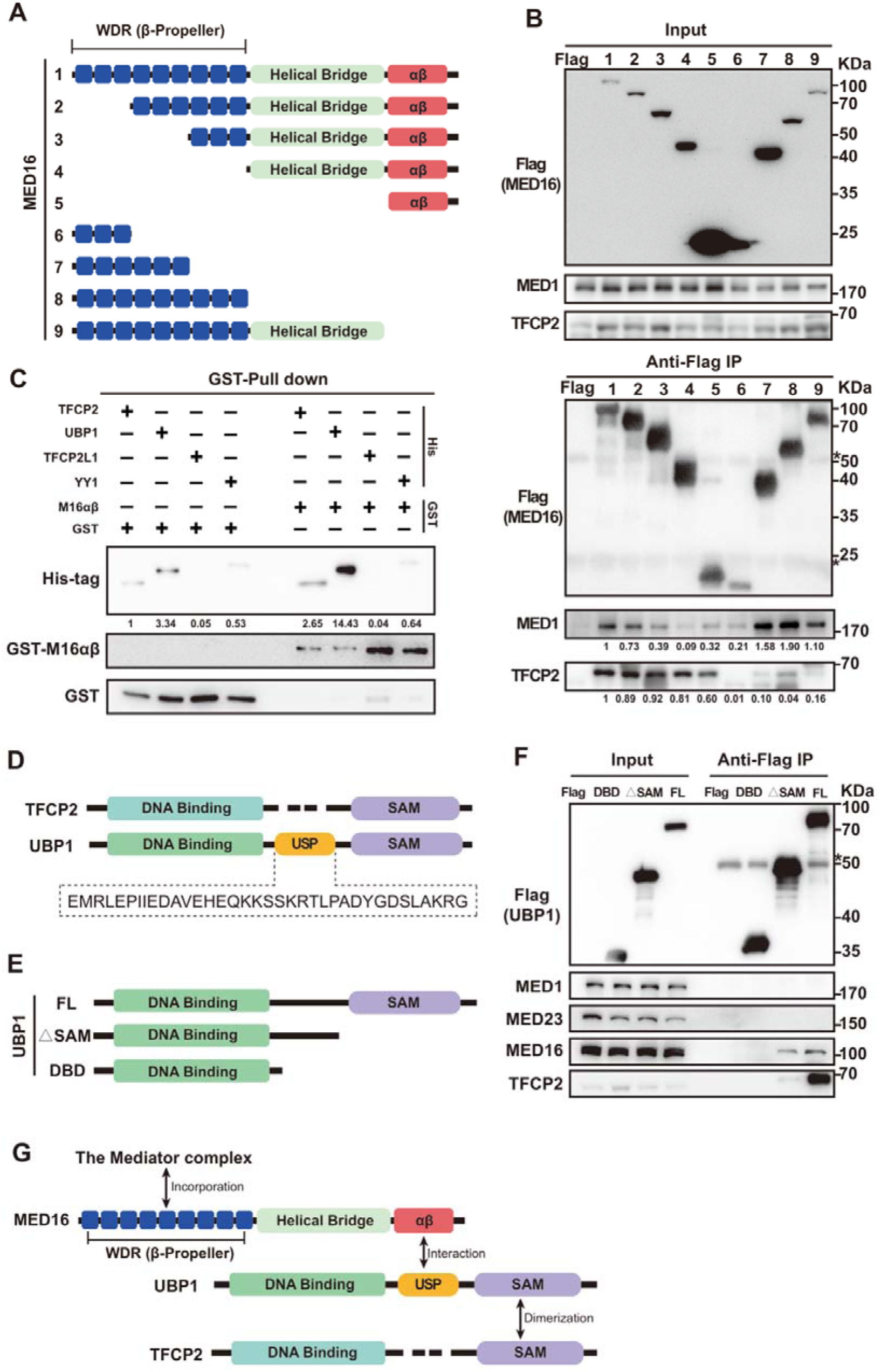
The N-terminal WDR domain of MED16 anchors it within the Mediator, while its C-terminal αβ-domain binds UBP1. (A) Schematic of Flag-MED16 deletion mutants. The WDR domains are shown as blue boxes. “1” represent full-length MED16 and “2” to “9” represent different truncations of MED16. All proteins were labeled with Flag-tag and expressed in 293T cell line as panel (B) indicated. (B) Domain mapping experiment of MED16. Flag-MED16 deletion mutants were coimmunoprecipitated with endogenous MED1 and TFCP2 using anti-Flag antibody. Asterisks indicate the IgG heavy and light chains. (C) Purified GST and GST-MED16 αβ-domain proteins were incubated with 6×His-tag TFCP2, UBP1, TFCP2L1, and YY1 at 40. After washing the interaction protein were eluted by boiling and immunoblotted with antibodies against GST-tag and His-tag. (D) Amino acids sequence analysis of UBP1 and TFCP2 protein, a 36 amino acids UBP1-specific sequence (USP) as shown below. (E) Schematic of Flag-UBP1 deletion mutants. FL represents full-length UBP1, △SAM represents SAM motif truncated UBP1, and DBD represents UBP1 DNA binding domain. (F) Domain mapping experiment of UBP1. Flag-UBP1 deletion mutants were coimmunoprecipitated with endogenous MED1, MED23, MED16, and TFCP2. Asterisks indicate the IgG heavy and light chains. (G) Schematic summary of the domain mapping results. The N-terminal WDR domain of MED16 mediates its integration into the Mediator complex, while the C-terminal αβ-domain binds UBP1. UBP1 interacts with MED16 through its UBP1-specific sequence (USP) and forms a heterodimer with TFCP2 via its SAM motif.

GST pull-down assays using purified recombinant proteins further validated direct binding. The GST-tagged MED16 αβ-domain exhibited robust direct interaction with His-tagged UBP1, with minimal binding to TFCP2, TFCP2L1, or YY1 (Fig 2C). Given the high sequence similarity between UBP1 and TFCP2, we further identified a unique 36-amino-acid specific peptide (USP) in UBP1 located between the DNA-binding domain and SAM motif (Fig 2D). Domain-mapping assays confirmed that the USP sequence is indispensable for MED16 binding, whereas the SAM motif mediates UBP1-TFCP2 dimerization (Fig 2E, 2F).

Together, these results establish that MED16 bridges UBP1/TFCP2 dimer to the Mediator complex. Specifically, the MED16 N-terminal WDR domain is responsible for anchoring MED16 to the Mediator, while MED16 C-terminal αβ-domain binds UBP1, with the USP sequence serving as the key recognition region for this interaction.

### MED16 acts through the Mediator complex to support UBP1-activated transcription

Because the interaction between transcription factors and Mediator subunits is critical for target gene activation, we analyzed if Mediator MED16 is required for UBP1-driven transcription activation using reporter gene assay (Fig 3A). In the co-transfection assay, the full-length UBP1 significantly promoted transcriptional activation. However, the DNA-binding domain of UBP1 (DBD, 1-274aa) alone was unable to activate the reporter. The truncated UBP1 mutant (ΔSAM, 1-369aa), which contains the MED16 binding motif but lacks the SAM motif for dimerization with TFCP2, exhibited about half the transcriptional activation when compared to the full-length UBP1 (FL) (Fig 3B).

**Figure 3.**
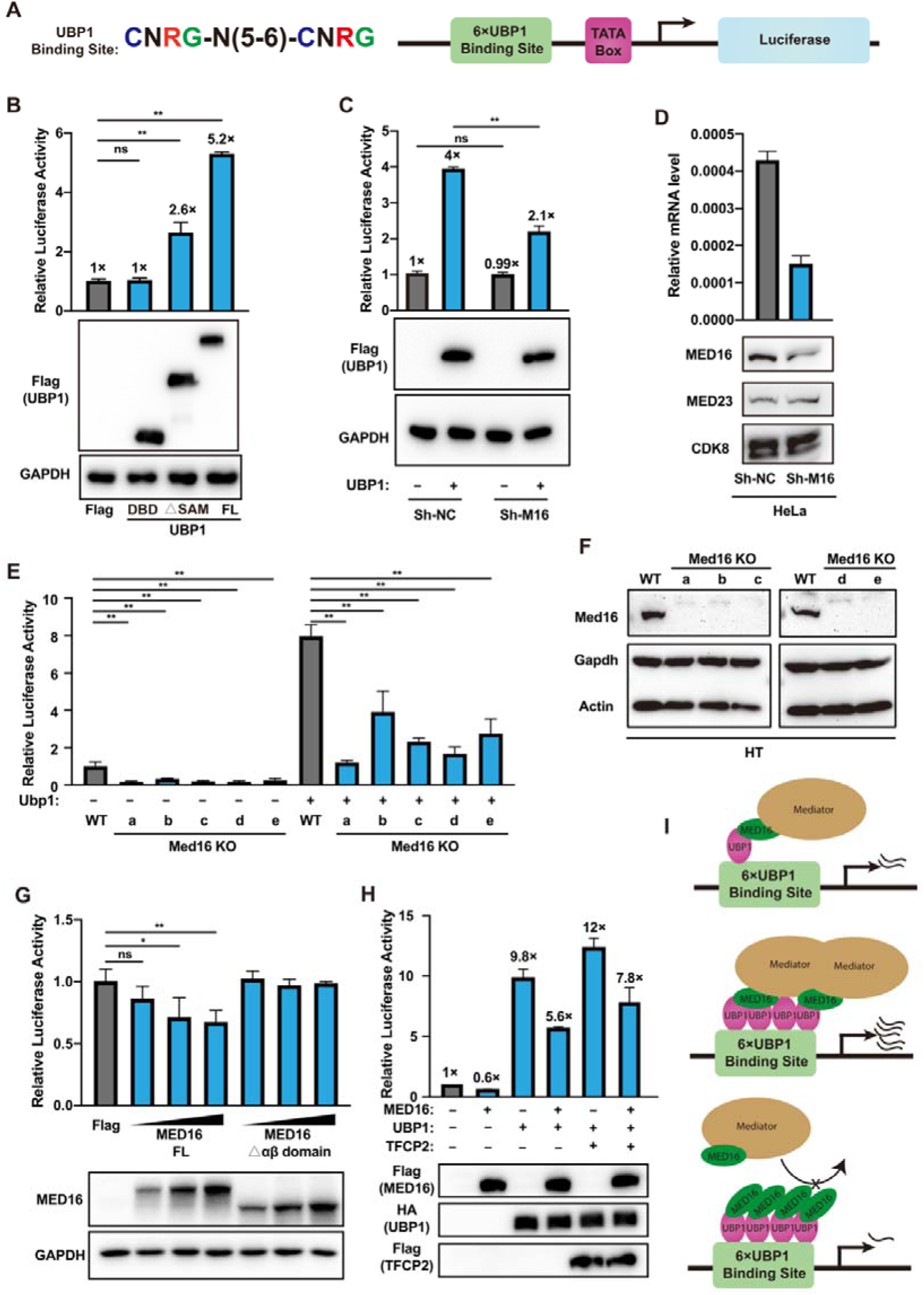
MED16 acts through the Mediator complex to support UBP1-mediated transcriptional activation. (A) Schematic of the UBP1 luciferase reporter construct. (B) Reporter activity in HeLa cells co-transfected with the reporter and 300 ng of plasmid expressing full-length UBP1 (FL), the SAM motif-deleted mutant (ΔSAM), or the DNA binding domain alone (DBD). (C) Reporter activity in control (sh-NC) and MED16-knockdown (sh-MED16) HeLa cells with or without UBP1 overexpression. (D) Knockdown efficiency of sh-MED16 confirmed by qPCR and Western blotting. (E) Reporter activity in wild-type and MED16 knockout (MED16 KO) mouse pancreatic cancer HT cells with or without UBP1 overexpression. (F) MED16 knockout efficiency confirmed by Western blotting in five independent frameshift clones. (G) Dose-dependent effect of wild-type MED16 or MED16Δαβ on UBP1 reporter activity. (H) Combinatorial effects of MED16, UBP1, and TFCP2 overexpression on UBP1 reporter activity. (I) Schematic model of UBP1-mediated transcriptional activation. (Top) UBP1 recruits the Mediator complex to the promoter through MED16 to activate transcription. (Middle) Overexpression of UBP1 enhances Mediator complex recruitment, leading to more active transcription. (Bottom) Overexpression of MED16 sequesters UBP1 away from the endogenous Mediator complex (squelching), thereby reducing Mediator recruitment and inhibiting transcription. Firefly luciferase activity was normalized to Renilla luciferase activity. Data are presented as means ± SD from n = 3 independent experiments. *: *p* < 0.05, **: *p*-value<0.01 and “ns”: no significant.

To further confirm that MED16 is required for the transcriptional activity of UBP1, a HeLa cell line with MED16 knockdown was generated using viral-mediated shRNA targeting MED16, along with a negative control cell line (sh-NC) (Fig 3D). The UBP1-responsive reporter showed similar baseline of transcriptional activity in the two cell lines. The reporter was activated about 4-fold by overexpressing UBP1 in the negative control cell line, whereas knockdown of MED16 attenuated the activation to approximately two-fold (Fig 3C).

The coding sequences of human *MED16* and Mouse *Med16* share about 90% identity. Thus, we further established a *Med16* knockout cell line generated using CRISPR-Cas9 gene editing method in a mouse pancreatic cancer cell line called HT. Five *Med16* knockout clones (ko16a, ko16b, ko16c, ko16d, & ko16e), with different frameshift mutations were selected and verified by sequencing and Western-blotting (Fig 3F). All *Med16*-KO clones displayed 2-7-fold reduced reporter activity under ectopic UBP1 activation when compared to the wildtype HT cells, and *Med16* KO exhibited decreased basal level of UBP1-reporter activity without UBP1 overexpression (Fig 3E and 3F). These results confirmed the specific requirement of MED16 for UBP1-mediated transcriptional activation.

To determine whether UBP1 requires the intact Mediator complex to activate transcription, we performed squelching experiments, based on the principle that overexpression of a Mediator subunit can sequester a transcription factor away from interacting with the endogenous Mediator complex, thereby inhibiting transcription 50-53). Wild-type MED16 repressed UBP1 reporter activity in a dose-dependent manner, consistent with this squelching effect. In contrast, increasing dose of MED16Δαβ, which cannot bind UBP1, had no inhibitory effect (Fig 3G). The same pattern was observed when UBP1 was co-expressed with TFCP2 (Fig 3H). These results indicate that UBP1-mediated transcriptional activation requires the intact Mediator complex, not MED16 acting alone.

### Analysis of the UBP1 and MED16 co-regulated transcriptome

To characterize MED16-UBP1-regulated transcription at the genomic scale, we performed RNA-seq analyses in control and MED16-knockdown HeLa cells, combining with or without UBP1 overexpression. A total of 217 genes were significantly activated by UBP1 overexpression, and 60 genes were repressed (fold change > 1.5, p < 0.05). Among the UBP1-activated 217 genes, 198 were profoundly downregulated upon MED16 depletion, supporting that Mediator MED16 is largely required for UBP1-dependent transcriptional activation (Fig 4A-4C).

**Figure 4.**
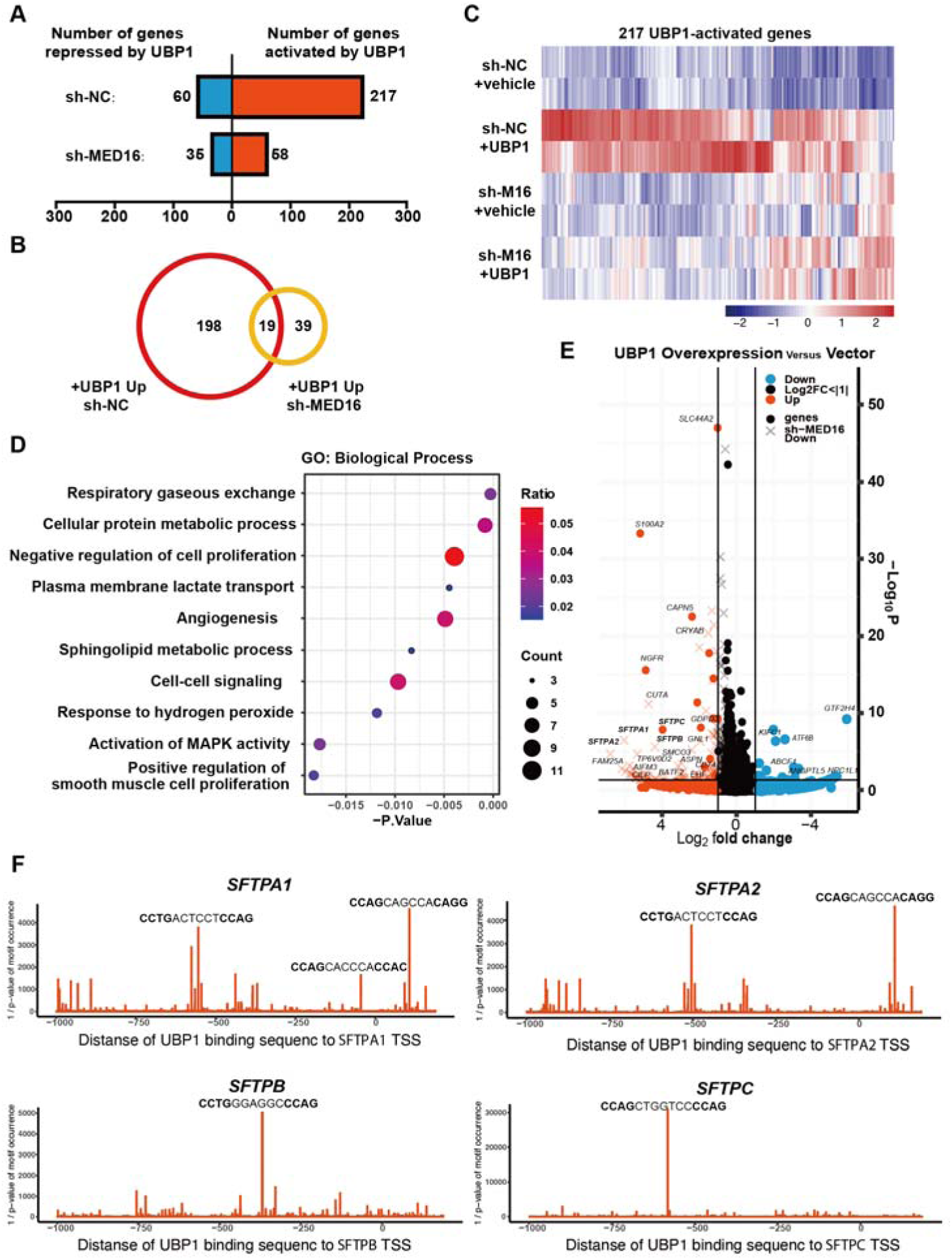
UBP1 and Mediator MED16 co-regulated transcriptome analysis. (A) The number of genes repressed or activated by UBP1 overexpression in the sh-NC and shMED16 HeLa cell line (FC > |1.5|, p ≤ 0.05). (B) Venn diagram of UBP1-activated genes in the sh-NC HeLa cell line and the shMED16 HeLa cell line (FC > |1.5|, p ≤ 0.05). 198 genes which were activated by UBP1 overexpression but abolished by sh-MED16 were defined as UBP1 and MED16 co-regulated genes. (C) Heatmap showing the RNA-seq analysis of 217 UBP1-activated genes in sh-MED16 and sh-NC cell line treated with or without UBP1 overexpression. UBP1-activated genes were defined in sh-NC cell line under UBP1 overexpression at fold change>|1.5|, p-value≤0.05. (D) GO analysis of 198 UBP1 and MED16 co-regulated genes. The top 10 biological process items were presented. (E) Volcano map analysis of differentially expressed genes in sh-NC HeLa cell line with or without UBP1 overexpression. Genes failed to be activated by UBP1 in the sh-MED16 HeLa cell line we’re indicated by “×”. (F) UBP1 motif occurrence analysis of *SFTPA1*, *SFTPA2*, *SFTPB* and *SFTPC*. Motif occurrence were showed within the promoter region of the genes (−1000 bp to 200 bp relative to the TSS).

To explore the potential biological processes that are co-regulated by MED16 and UBP1, 198 co-regulated genes by MED16 and UBP1 were subjected to gene ontology (GO) analysis (Fig 4D and 4E). Interestingly, GO analysis revealed that the genes involved in the respiratory gaseous exchange process were regulated by MED16 and UBP1, such as *SFTPA1*, *SFTPA2*, *SFTPB*, and *SFTPC* (Fig 4F). These genes encode pulmonary surfactant associated proteins that promote alveolar stability by lowering the surface tension at the air-liquid interface in the peripheral air spaces. They are normally only expressed in alveolar epithelial cells and silenced in HeLa cells. Interestingly, highly matched UBP1 motifs were identified in their promoters (Fig 4F), and their activation by UBP1 overexpression is associated with the presence of these motifs. In our analysis, angiogenesis is also enriched in GO analysis (Fig 4D), which is consistent with the previous report that UBP1 plays a critical role in the regulation of extraembryonic angiogenesis (29, 54). Collectively, these results demonstrated that UBP1 and MED16 may collaborate to activate a target array of endogenous genes expression.

### Mediator MED16 collaborates with UBP1 to inhibit HIV-1 transcription

Beyond their roles in transcriptional activation, UBP1 and TFCP2 have been reported to inhibit HIV-1 transcription through multiple mechanisms (32–34). We therefore asked whether MED16 participates in UBP1-mediated HIV-1 repression. To test this, we constructed an HIV-1 reporter driven by the viral core promoter (−78 to +60) upstream of luciferase (Fig 5A). Overexpression of MED16, UBP1 or YY1 reduced reporter activity, suggesting that MED16 behaves similarly to the known HIV-1 transcriptional repressors (Fig 5B). Overexpression of MED16 also inhibited the reporter in a dose-dependent manner (Fig 5E), whereas MED23 and MED24 had no effect (Fig 5C). Conversely, shRNA-mediated knockdown of MED16 modestly increased HIV-1 reporter activity (Fig 5D). These results indicate a specific role for MED16 in repressing HIV-1 transcription.

**Figure 5.**
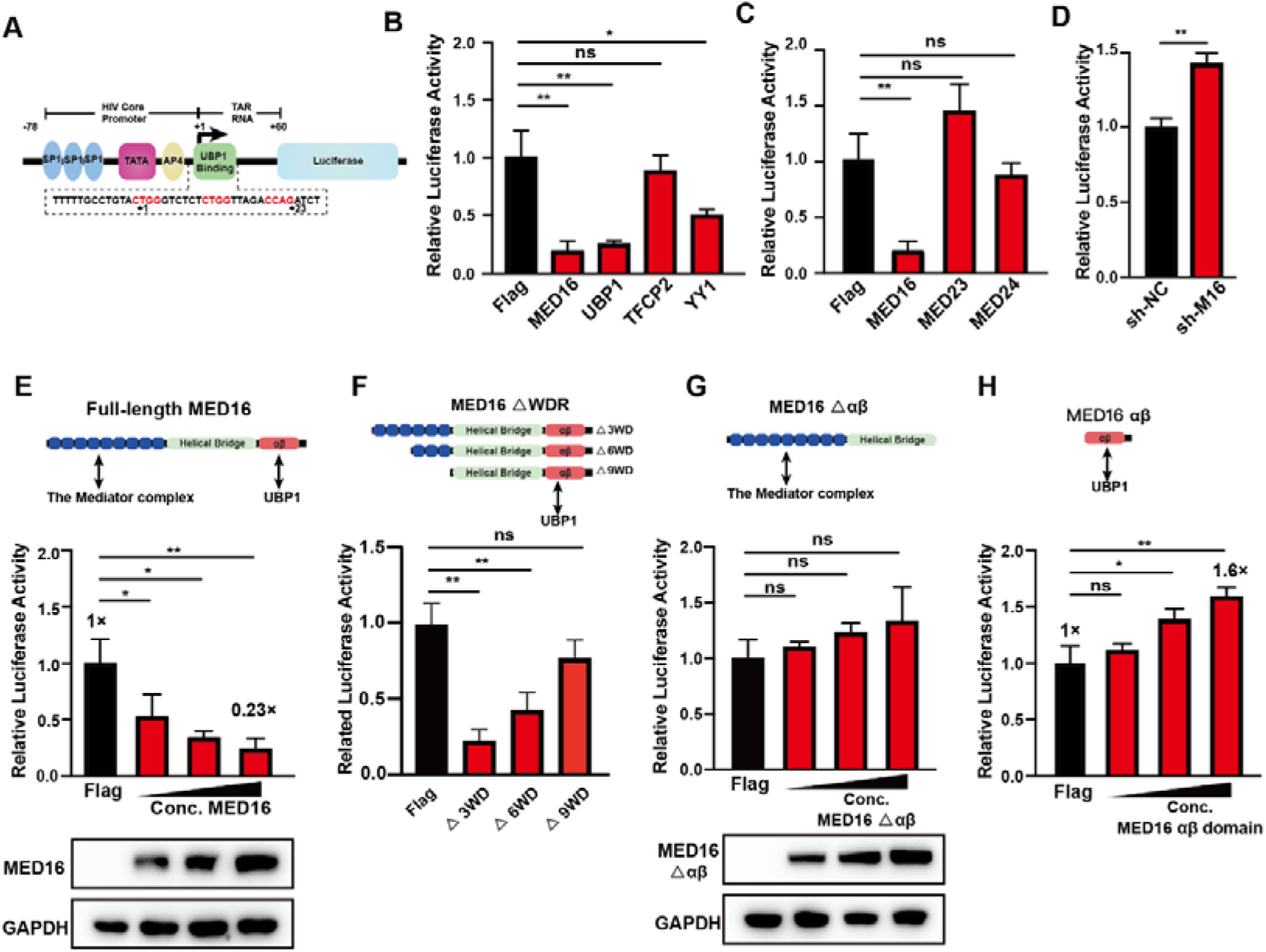
Mediator MED16 cooperates with UBP1 to inhibit HIV-1 reporter expression. (A) Schematic illustration of the HIV-1 reporter, with the HIV-1 core promoter and +1 to +60 TAR RNA sequence cloned upstream of the luciferase coding sequence (CDS). (B) The HIV-1 reporter was co-transfected with 1.5 μg of MED16, UBP1, TFCP2, YY1, or Flag-tag control plasmids in 293T cells. (C and D) Increasing amounts (0.7 μg, 1 μg, 1.5 μg) of MED16 or αβ-domain-deleted MED16 were transfected with the HIV-1 reporter. (E) The HIV-1 reporter was co-transfected with 1.5 μg of MED23, MED24, or Flag-tag control plasmids as additional controls. (F) Comparison of HIV-1 reporter luciferase activity between sh-NC and sh-MED16 293T cell lines. Firefly luciferase activity was normalized to Renilla luciferase activity. The luciferase expression levels between groups were compared using two-tail unpaired Student’s t-test, The data are presented as the means ± SDs from n = 3 independent experiment. * *p* < 0.05, ** *p*-value<0.01 and “ns”: no significant.

We next asked whether this repression requires the Mediator complex and the interaction with UBP1. The MED16ΔWDRs mutants, which cannot be integrated into the Mediator, failed to inhibit the HIV-1 reporter, indicating that MED16 must act through the Mediator complex to repress HIV-1 transcription (Fig 5F). A MED16 mutant lacking the C-terminal αβ-domain (MED16Δαβ), which cannot bind UBP1, also failed to inhibit the reporter (Fig 5G). Conversely, overexpression of the MED16 αβ-domain peptide, which competes with endogenous MED16 for UBP1 binding, de-repressed HIV-1 reporter activity (Fig H). Together, these results demonstrate that the functionality of MED16 in HIV-1 repression depends on both its integration into the Mediator complex and its interaction with UBP1.

### UBP1 binding position determines the MED16-dependent inhibition of HIV-1 transcription

In a previous study, UBP1 was shown to inhibit HIV-1 transcription by recruiting YY1 and HDAC1(34). To investigate if MED16 employs a similar mechanism to repress HIV-1 transcription, we transfected HeLa cells with an HIV-1 reporter and treated them with panobinostat, a pan-HDAC inhibitor. Western blot analysis showed increased acetylation of histone H3 (H3ac) upon panobinostat treatment (Fig S2A). However, this treatment did not reverse MED16-mediated inhibition of HIV-driven luciferase activity (Fig S2A). This result suggests that MED16 may not rely on HDAC-mediated histone deacetylation to inhibit HIV-1 transcription and alternative mechanisms may be involved in HIV-1 transcription repression.

We then asked whether the cis regulatory elements in the HIV-1 promoter are necessary for MED16-mediated inhibition. Firstly, we examined whether the transactivating response region (TAR) plays a role in MED16-mediated HIV-1 inhibition HIV-1 inhibition. TAR is transcribed into TAR RNA, which recruits the host P-TEFb-containing super elongation complex (SEC) (55–57) to mediate the activation of viral protein Tat (58). UBP1 has been reported to restrict HIV-1 transcription at the level of elongation (33). A TAR RNA mutation HIV-1 reporter was created by replacing TAR loop nucleotides 31-34 UGGG with CAAA, which reduces binding affinity of Tat:P-TEFb and abolishes the function of TAR RNA (59). We observed that MED16 inhibited both wild-type and TAR-mutant reporter’s transcription, while mutated TAR attenuated UBP1-mediated inhibition (Fig S2B and S2C). This result suggested that MED16-mediated inhibition of HIV-1 is independent of the TAR-mediated HIV-1 transcriptional elongation.

HIV-1 proximal promoter has a string of GC-box that is bound by SP1, and an AP4 binding site between TATA-box and TSS (60). We then examined whether these two transcription factors are involved in the MED16-mediated transcription repression. A reporter driven by 4×GC-box was co-transfected with the MED16 expression plasmid, but no significant inhibition was observed compared with co-transfecting with the control vector (Fig S2D), suggesting no cross-talk between MED16 and Sp1 (GC-box). Then we mutated the AP-4 binding motif CAGCTG to CAGTCG (Fig S2E) to abolish AP-4 binding (61). Overexpressing MED16 inhibited the luciferase activity, regardless AP-4 binding status, suggesting that MED16 doesn’t work with AP-4 to inhibit the HIV-1 transcription.

Lastly, we examined if the UBP1-MED16 inhibition of HIV-1 transcription is due to the UBP1 binding position at the transcription started site (TSS) (32, 33). Two chimera reporters, both driven by a chicken β*-ACTIN* promoter, were created with distinctive UBP1 binding site position. In the first reporter (Fig 6A), the native TSS of the chicken β*-ACTIN* promoter was replaced with the HIV-1 TSS, which contains a UBP1 binding site. In the second reporter (Fig 6B), a UBP1 binding site was placed upstream of the chicken β*-ACTIN* TSS. Various truncated mutants of MED16 and UBP1 were co-transfected with these reporters, respectively. In the case of the first chimera reporter, transcription was inhibited by full-length MED16 and MED16-bound UBP1 variants (ΔSAM and FL-UBP1), but not by UBP1-binding deficient MED16 (MED16Δαβ) or MED16-binding deficient UBP1 (UBP1-DBD) (Fig 6A). Conversely, transcription of the second chimera reporter was not inhibited by any MED16 or UBP1 variants (Fig 6B). Additionally, comparison of transcription activities between the two reporters revealed that the first chimera exhibited approximately 50% lower transcription activity than the second, possibly due to the endogenous UBP1 activity (Fig 6C). These findings suggest that the UBP1 binding at the TSS seems to be key factor for MED16-mediated transcriptional repression

**Figure 6.**
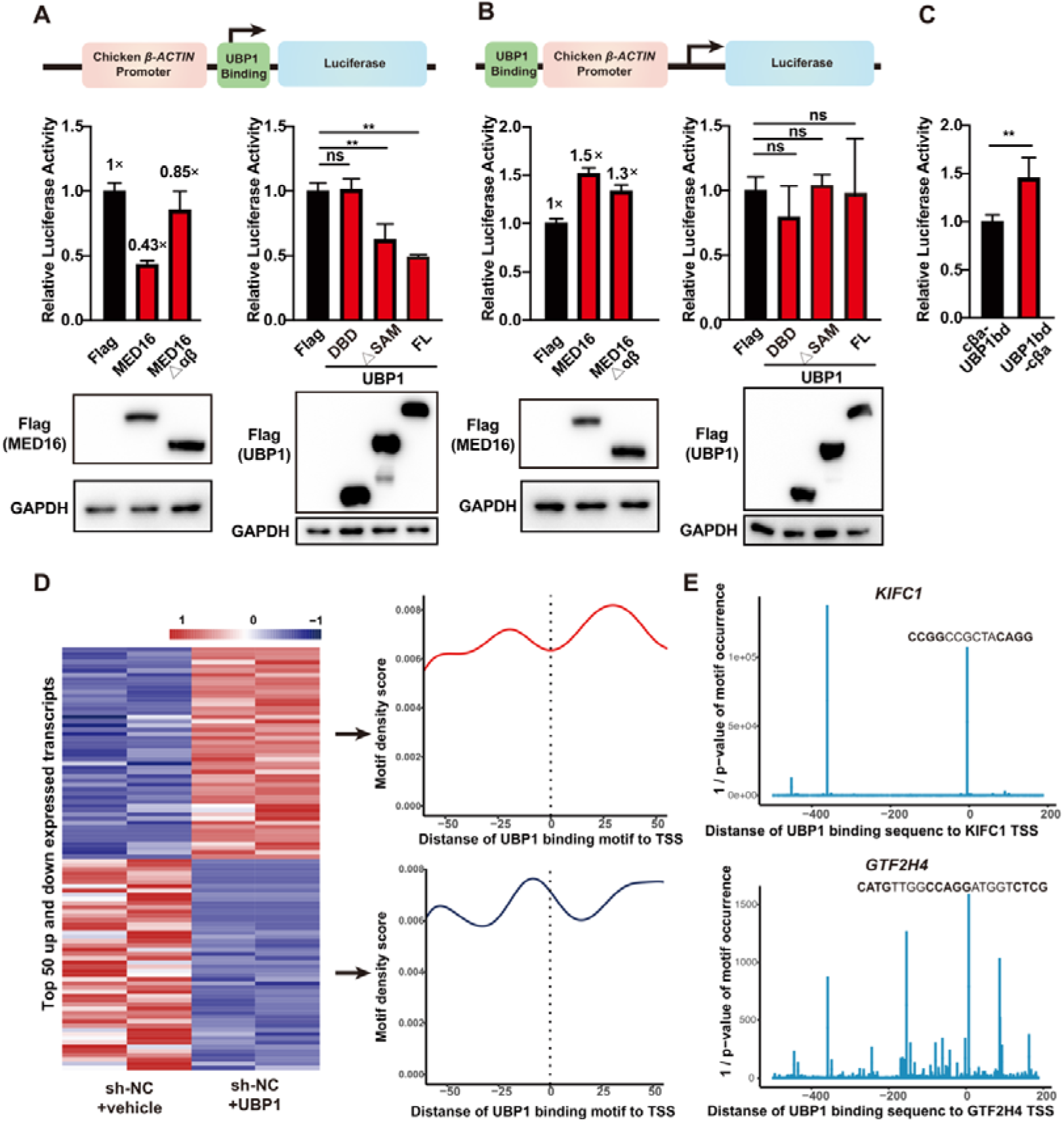
The position of the UBP1 binding site relative to the TSS dictates whether MED16-UBP1 activates or represses transcription. Schematic showing a chimera reporter (A) driven by the Chicken β*-ACTIN* promoter, followed by an HIV-1 UBP1 binding site that also serves as the HIV-1 transcription start site and another chimera reporter (B) driven by the Chicken β*-ACTIN* promoter, a UBP1 binding site modify from HIV-1 TSS placed upstream of it. 1 μg of different truncation of MED16 or UBP1 plasmid was co-transfected with the reporter. The Luciferase activity comparison between two chimera reporters were shown in (C). “cβa-UBP1 bd” indicate the reporter that panel A described, and “UBP1 bd-cβa” indicate the reporter that panel B described. Firefly luciferase activity was normalized to Renilla luciferase activity. The luciferase expression levels between groups were compared using two-tail unpaired Student’s t-test, The data are presented as the means ± SDs from n = 3 independent experiment. * *p* < 0.05, ** *p*-value<0.01 and “ns”: no significant. UBP1 motif enrichment analysis of top 50 upregulated or top 50 downregulated transcripts induced by UBP1 were shown in (D). The transcripts changes levels are shown on the left as a heatmap, and average profile plots of UBP1 motif density are shown on the right. UBP1 motif occurrence analysis of UBP1 downregulated gene *KIFC1* and *GTF2H4* were shown in (E). Motif occurrence were showed within the promoter region of the genes (−500 bp to 200 bp relative to the TSS).

We then looked into whether the binding positional effect of UBP1 also impacted on the UBP1-tageted gene expression across the human genome. The correlation between the relative binding position of UBP1 to TSS and gene transcription levels was analyzed using the gene profiling data (Fig 4). We selected the top 50 upregulated and top 50 downregulated transcripts under UBP1 overexpression and analyzed the enrichment of the UBP1 motif around their promoters (Fig 6D). Notably, UBP1-activated transcripts showed significant enrichment of the UBP1 motif upstream and downstream of the TSS, while UBP1-repressed transcripts exhibited motif enrichment specifically overlapping the TSS (Fig 6D). For example, two UBP1-inhibited gene: *KIFC1* and *GTF2H4*, were found to have their TSS overlapping with UBP1 binding motif (Fig 4F and 6E). Together with the chimeric reporter experiments, these data suggest that UBP1 can act as either a transcriptional activator or repressor, with its binding position relative to the TSS as a determinant of this two-way regulatory outcome.

### UBP1-MED16 interrupts HIV-1 transcription initiation and sustains HIV-1 latency

We wondered how UBP1-MED16 binding within the proximal promoter might influence the HIV-1 transcription. To directly investigate how MED16-UBP1 binding at the TSS represses HIV-1 transcription, we performed in vitro immobilized template transcription assays using nuclear extracts from wild-type, Med16-knockout, and Med16-overexpressing HT cells (Fig 7A). During pre-initiation complex (PIC) assembly (without NTPs addition), MED16 overexpression reduced the recruitment of CDK8, MED12, MED1, and MED23 to the HIV-1 promoter. Upon transcription initiation by NTPs addition, Pol II Ser5 phosphorylation was markedly diminished, while the retention of TFIIB, a GTF that dissociates after initiation, was prolonged (Fig 7B). Functionally, MED16 overexpression reduced RNA synthesis of the viral template by approximately 45% (Fig 7C). These results suggest that MED16 overexpression disrupts the PIC formation for HIV-1 transcription initiation, leading to a reduction in RNA production.

**Figure 7.**
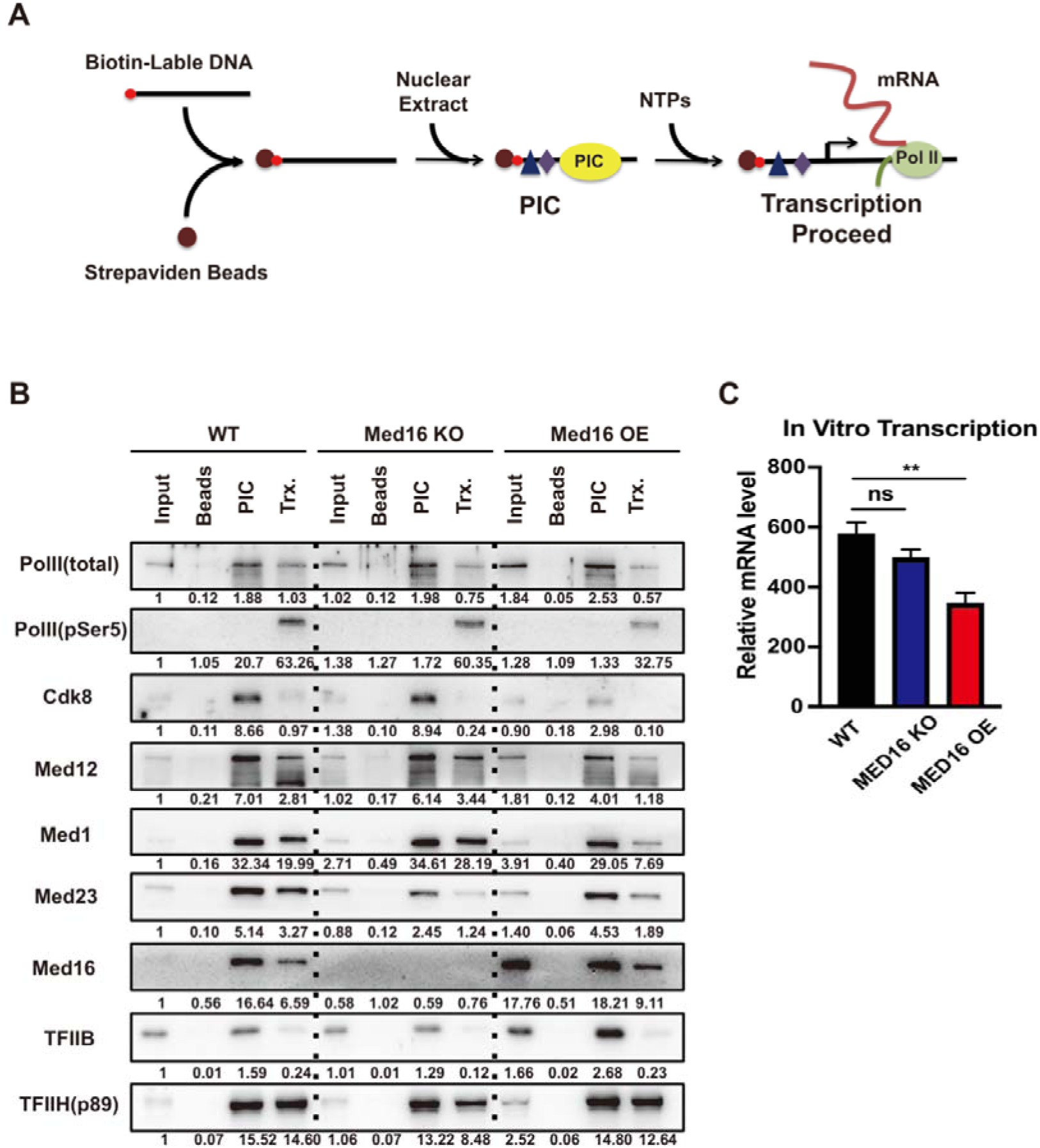
MED16 interferes with HIV-1 PIC assembly at the TSS via its interaction with UBP1. (A) Schematic of biotin-labeled immobilized template assay. (B) Immobilized template assay in wild-type, Med16 knockout and Med16 overexpression HT cell line. The binding proteins in individual steps were collected for immunoblotting. Beads: the streptavidin beads incubated with NE as a negative control; PIC: the pre-initiation step of transcription; Trx.: transcription initiation from the PIC step by adding NTPs. (C) qPCR quantification of generated RNA from different NE. The mRNA expression levels were compared between groups using two-tail unpaired Student’s t-test, The values are presented as the means ± SDs from n = 3 independent experiment. ** *p*-value<0.01 and “ns”: no significant.

Because the MED16-UBP1 interaction inhibits HIV-1 transcription (Figure 5), we went on exploring whether it might also play a role in maintaining HIV-1 latency in CD4^+^ T cell line. For this purpose, we used the J-Lat 10.6 cell line, a Jurkat-based cell line infected with a pseudotyped HIV-1 strain (HIV/R7/E-/GFP) (62). This cell line harbors the integrated HIV-1 copy in the second intron of the *SEC16* gene (63) and the integrated HIV-1 copy remains silenced under normal conditions. HIV-1 latency reversal can be monitored by GFP expression that is controlled by viral promoter and serves as a marker for viral activation (Fig 8A). To induce viral expression, we treated cells with JQ1, a BET bromodomain inhibitor that reverses HIV-1 latency primarily by antagonizing BRD4-L-mediated suppression of the Tat-P-TEFb/SEC interaction (64, 65), while displacement of the BRD4-S isoform from the viral promoter may further contribute to this effect (66, 67).

**Figure 8.**
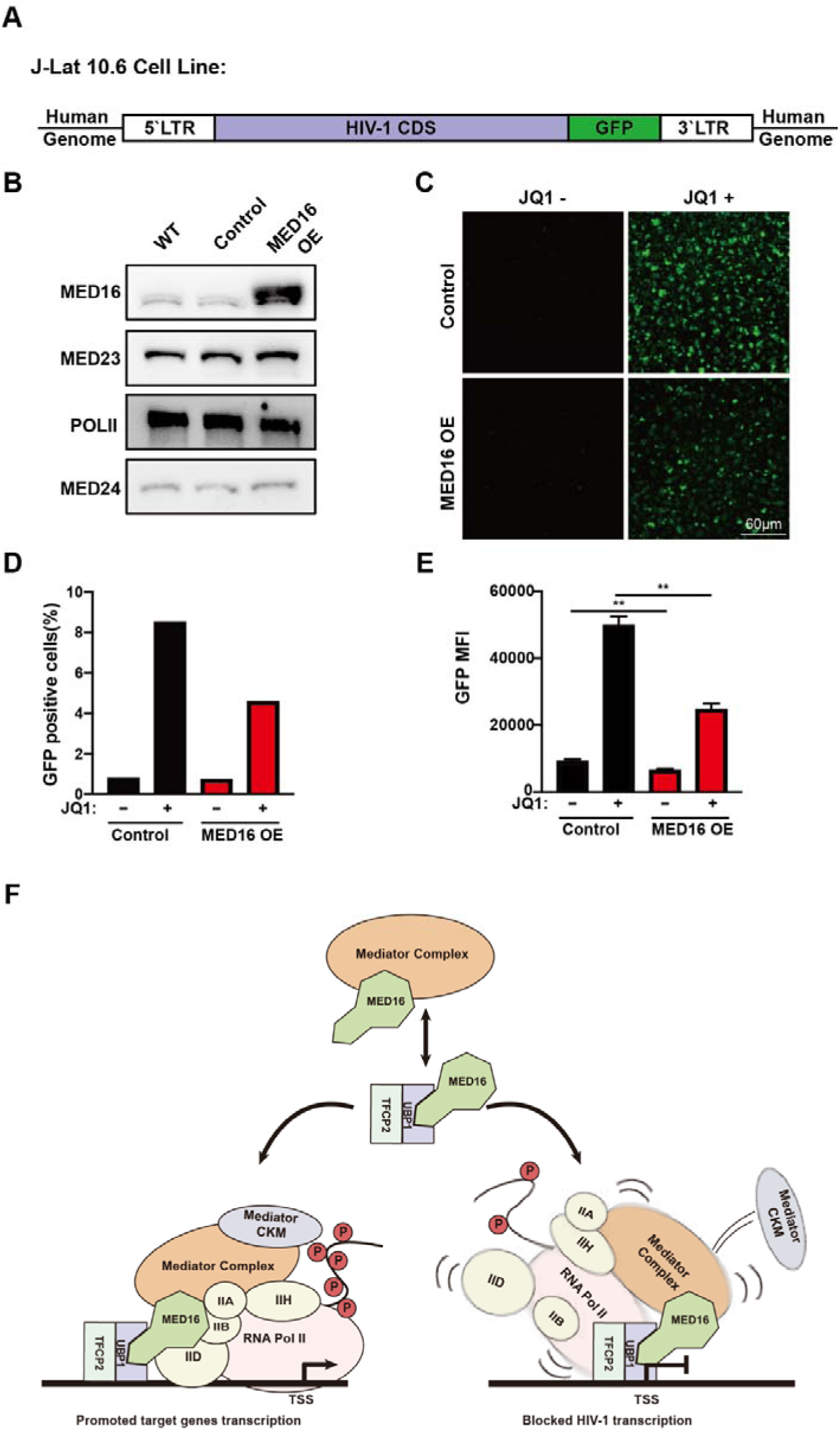
Mediator MED16 prohibited PIC formation of HIV-1 transcription. (A) Schematic of the J-Lat 10.6 cell line, a Jurkat-derived CD4+ T cell model harboring an integrated latent HIV-1 provirus (HIV/R7/E-/GFP) in the second intron of the *SEC16* gene. Under normal conditions, the provirus remains silenced, and latency reversal can be monitored by GFP expression driven by the viral promoter. (B) Western blot analysis confirming MED16 overexpression in MED16 OE J-Lat 10.6 cells compared with the control cell line. (C) Representative fluorescence microscopy images of MED16 OE and Control J-Lat 10.6 cells treated with or without 5 μM JQ1 for 48 hours. (D) Percentage of GFP-positive cells quantified by FACS. (E) Mean fluorescence intensity (MFI) measured using the BioTek Synergy 2 system. (F) Schematic model of MED16-mediated transcriptional regulation. MED16 uses its N-terminal WDR domain to anchor within the Mediator complex and its C-terminal αβ-domain to bind UBP1. When the UBP1 binding motif is located upstream of the TSS, UBP1 recruits the Mediator complex through MED16 to activate transcription. When the motif overlaps the TSS, as in the HIV-1 promoter, this interaction instead disrupts PIC assembly, leading to transcriptional repression and the maintenance of HIV-1 latency. The values are presented as the means ± SDs from n = 3 independent experiment. ** p-value<0.01

We generated J-Lat 10.6 cell lines with retroviral MED16 expression and a control line infected with the MSCV (Murine Stem Cell Virus) retrovirus (Fig 8B). Prior to JQ1 treatment, both cell lines showed minimal GFP expression, confirming that MED16 manipulation alone did not alter HIV-1 latency (Fig 8C). After treatment with 5μM JQ1 for 48h, the HIV-LTR-driven GFP expression is observed in both cell lines, but is reduced in the MED16 OE cell line, with a 40% decrease in the percentage of GFP-positive cells percentage and 0.54-fold reduction in MFI (mean fluorescence intensity) compared to the control cell line (Fig 8C, 8D, and 8E). These findings suggest that MED16 not only inhibits HIV-1 transcription but also plays a important role in sustaining HIV-1 latency in CD4^+^ T cells.

## DISCUSSION

In this study we identified that the UBP1-TFCP2 heterodimer interacts with Mediator subunit MED16. This interaction activates an array of endogenous gene expression and inhibits HIV-1 transcription.

Using gel-filtration chromatography and IP-MS, we first observed that transcription factor UBP1 and TFCP2 are associated with MED16 from the biochemically resolved fractions (Fig 1D, 1E and 1F). Although these transcription factors were initially found in MED16-containing fractions outside the core Mediator, our functional experiments demonstrate that they rely on the Mediator complex for transcriptional regulation. In the context of gene activation, squelching experiments showed that overexpressing wild-type MED16 inhibited UBP1-driven reporter activity, whereas the MED16△WDR and MED16△αβ mutants, which cannot either incorporate into Mediator or bind to UBP1, had no inhibitory effect (Fig 3G, 3H, 5G). This indicates that UBP1-mediated activation requires the intact Mediator complex, not MED16 alone. In the context of gene repression, MED16△WDR, a MED16 mutant lacking the N-terminal WDR domain, which cannot integrate into the Mediator, failed to inhibit the HIV-1 reporter (Fig 5F), demonstrating that UBP1-mediated repression also requires the whole Mediator complex. Thus, in either activating cellular genes or repressing HIV-1 transcription, UBP1 function by recruiting the Mediator complex through MED16 (Fig 8F). In this model, MED16 acts as a bridging subunit whose N-terminal WDR domain integrates it into the Mediator and whose C-terminal αβ-domain directly binds UBP1, thereby physically linking the transcription factor to the coactivator complex.

Previous studies have demonstrated UBP1’s role in inhibiting HIV-1 transcription through various mechanisms, including epigenetic and elongational regulation. Our investigation reveals a novel mechanism where the Mediator MED16-UBP1 interaction inhibits HIV-1 transcription initiation by compromising PIC formation. This finding highlights the importance of UBP1 binding site positioning, a concept recently underscored by Duttke and colleagues, who demonstrated that the spatial configuration of TF binding sites dictates transcriptional outcomes (68). Our work also extends this principle: UBP1 activates transcription when binding upstream or downstream of the TSS, but represses it when binding directly at the TSS. Additionally, we identified TFCP2 and the known HIV-1 silencing factor YY1 (34) in the MED16-containing fractions, although their precise roles in our mechanistic model remain to be determined.

Targeted inhibition of HIV-1 transcription, particularly through enhancing the silencing activity of UBP1-MED16 at the viral transcription start site (TSS), presents a potential tool for HIV-1 therapy. Many current approaches focus on inhibiting key regulatory proteins crucial for HIV-1 transcription (69–71). By strengthening the UBP1-MED16 interaction or increasing MED16’s affinity for the HIV-1 TSS, it may be possible to achieve more effective transcriptional silencing, which could reduce viral expression over prolonged periods. In the future, small molecule drug screening can be conducted targeting the enhancement of this interaction, and the obtained small molecules could be used in combination with existing antiretroviral therapies to enhance therapeutic efficacy (72, 73).

Interestingly, a small peptide fragment (αβ-domain) of MED16 has the opposing effect of activating HIV-1 transcription while potentially blocking the endogenous UBP1-MED16 interaction (Fig 5H). By manipulating the expressed form of MED16, we may be able to create a multi-faceted therapeutic tool that enables flexible regulation of HIV-1 transcription based on specific therapeutic needs. For instance, expressing the full-length MED16 could enhance the efficacy of antiretroviral therapies (ART), while expressing the MED16 αβ-domain peptide could support “shock-and-kill” strategies (74) by reactivating latent HIV-1, thereby enabling immune clearance or targeted antiretroviral intervention in previously hidden reservoirs. Together, these dual capabilities of MED16 may provide multiple strategies in the HIV-1 intervention.

## Method

### Plasmids

The human *MED16*, *YY1*, *TFCP2*, *TFCP2L1* cDNAs were obtained from BRICS (Bio-Research Innovation center SUZHOU). The human *UBP1* and mouse *Med16* cDNAs were cloned from HeLa cells and HT cells respectively. Then full-length cDNA and truncations were cloned into 3×Flag-CMV-10 (Sigma-Aldrich-Aldrich) and pET-28b plasmids. The HIV-1 core promoter sequence were synthesized directly and clone into pGL3-Basic plasmid. All mutations on plasmid were generated by using FASR Site-Directed Mutagenesis Kit (Tiangen; Beijing; China). siMED16 and Ctrl shRNA oligonucleotides were cloned into pSiren-RetroQ (Clonetech).

### Cell culture

HeLa and 293T cells were maintained in Dulbeccòs modified Eaglès medium (DMEM) containing 10% (v/v) FBS and 1% P/S. The HT cell line used in this study was an immortalized pancreatic cancer cell line derived from a KRASG12D P53+/-mouse and was generous provided by professor Xiaofei Yu in Fudan University. J-Lat 10.6 cell line was generous provided by professor Huanzhang Zhu in Fudan University. HT and J-Lat 10.6 cell were maintained in RPMI-1640 medium containing 10% (v/v) FBS and 1% P/S. All cells were maintained in the incubator with 5% CO2 at 37□.

### Western blot and real-time PCR assays

Method for Western blot and real-time PCR assay have been described previously (75). The primers for real-time PCR are listed in Table S1. Antibody for Western blot include the following MED23 (Abcam, ab200351), MED1 (Bethyl Lab, A300-793A), MED24 (Bethyl, A301-472A), MED6 (Santa Cruz, sc-9434), MED12 (Bethyl Lab, A300-774A), MED16 (Bethyl Lab, A303-668A), CDK8 (Abcam, ab115155), UBP1 (Protientech, 67318-1-Ig), TFCP2 (Proteintech, 15203-1-AP), YY1 (Proteintech, 66281-1-Ig), P300 (Abcam, ab14984), HDAC1 (Santa Cruz, sc-7872), PolII (Abcam, ab816), PolII-pSer5 (Millipore, 04-1572), TFIIB (Santa Cruz, sc-274D), TFIIH (Santa Cruz, sc-293), H3 (Abcam, ab1791), H3ac (Abcam, ab4729), β-Actin (Proteintech, 66009-1-Ig), GAPDH (Proteintech, 60004-1-Ig), Flag-tag (Sigma-Aldrich, F1804), His-tag (Proteintech, 66005-1-Ig), GST-tag (Invitrogen, 10004D).

### Nuclear extract Preparation and gel-filtration chromatography assay

Preparation of un-dialyzed HeLa cell nuclear extract was described previously (76). Gel-filtration molecular weight markers (Sigma-Aldrich, MWGF1000) were loaded onto a Superose 6 10/300GL column to determine the void volume (Blue dextran ∼2000 kDa) and elution volume of protein standards (Thyoglobulin 669 kDa, Apoferritin 443kDa, Alcohol Dehydrogenase 150 kDa, Albumin 66 kDa). HeLa nuclear extract was applied to the Superose 6 10/300GL column and then was run in Buffer D (20 mM HEPES (pH 7.9), 300 mM KCl, 10% glycerol, 0.2 mM EDTA, 10 mM β-mercaptoethanol, and 1 mM PMSF). Fractions of 500µl were collected from the column and precipitated using trichloroacetic acid (TCA). Every third fraction was used for Western blot assay.

### Co-immunoprecipitation (Co-IP) assay

For transient co-transfection, 293T cells plated in 10 cm dishes (90% confluency) were transfected with 10 μg of each plasmid using Lipofectamine 2000 (Invitrogen) according to the manufacturer’s protocol. After 48 h, the cells were harvested and lysed in lysis buffer (20mM HEPES pH7.5, 150mM NaCl, 1mM EDTA, 2.5mM EGTA, 0.3% Tween-20) with protease inhibitors. Lysates were subjected to immunoprecipitation with anti-Flag M2 beads (Sigma-Aldrich, A2220) and incubated at 4□ overnight, following by washing in lysis buffer three times. Then 50 μl 1× SDS loading buffer was added to the beads and boiled at 990 for 10 min for Western blot analysis with the indicated antibodies.

For endogenous Co-IP, 293T cells were seeded in 10 cm dishes and cultured under appropriate conditions. At nearly 80% confluence, the cells were harvested and lysed in lysis buffer with protease inhibitors. Lysate was centrifuged at 4□, 13200 rpm for 10 minutes, and the supernatant was incubated with 4μg antibodies to MED16 (Bethyl, A303-668A), CDK8 (CST, 17396s), MED1 (Bethyl, A300-793A), MED12 (Bethyl, A300-774A) overnight. Then, 20 μl of Dynabeads Protein G (Invitrogen) was added and incubated for 2 hours at 4□. The beads were then washed with lysis buffer (with 0%, 5%, 10% 1,6-hexanediol) three times and boiled with SDS loading buffer for Western blot assay with indicated antibodies.

As for the Co-IP experiment in HeLa nuclear extracts (NE) or gel-filtration fractions, the HeLa NE was first diluted with an equal volume of D300 buffer containing 20 mM HEPES (pH 7.9), 300 mM KCl, 10% glycerol, 0.2 mM EDTA, 10 mM β-mercaptoethanol, and 1 mM PMSF. Next, 4 μg of antibody was added directly to the diluted HeLa NE or gel-filtration fractions, followed by incubation at 4°C overnight. After the overnight incubation, 20 μL of Dynabeads Protein G (Invitrogen) was added and incubated for an additional 2 hours at 4°C. The beads were then washed three times with D300 buffer and boiled in SDS loading buffer for Western blot analysis with the specified antibodies.

### IP-MS

After immunoprecipitation, the protein samples were separated by SDS0PAGE, running at 80 V for about 15 minutes. The electrophoresis was halted once the bromophenol blue dye front had traveled 0.5 cm into the resolving gel. The gel was then stained with Coomassie Brilliant Blue R250 to visualize protein bands, which were subsequently excised and diced into small fragments (1 mm³). These gel fragments underwent sequential washes: twice with a 50% acetonitrile (ACN) solution in 50 mM ammonium bicarbonate (NH_4_HCO_3_), followed by three washes with 10 mM NH_4_HCO_3_ and one wash with pure ACN to remove salts and detergents.

After dehydration with 100% ACN, the gel pieces were treated with 10 mM dithiothreitol (DTT) for 1 hour at 37 °C to reduce disulfide bonds. The DTT solution was then removed, and the gel fragments were dehydrated again with ACN. Next, 50 mM iodoacetamide (IAM) was added to alkylate free thiol groups, with incubation for 30 minutes in the dark. The gel pieces were washed three times with 10 mM NH_4_HCO_3_ and once with ACN before final dehydration.

For enzymatic digestion, the gel fragments were incubated overnight at 37 °C with trypsin (3 ng/μl in 10 mM NH_4_HCO_3_). The resulting peptides were extracted using a gradient ACN elution and analyzed by liquid chromatography-tandem mass spectrometry (LC-MS/MS) on an Orbitrap Fusion™ Lumos™ system.

### IP-MS data analysis

IP-MS was performed using antibodies against MED16, MED1, MED12, and CDK8, each with three biological replicates. For each bait, protein abundance was normalized to the total intensity of all identified Mediator subunits in the same sample. This normalized abundance was expressed as parts per million (PPM), correcting for differences in immunoprecipitation efficiency across baits.

For each protein *g* in sample *s*, we first calculated the area-based abundance in parts per million:

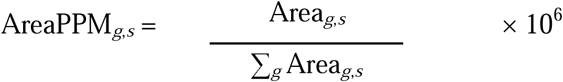

This value was then log2-transformed:

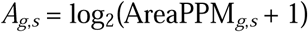

To correct for differences in immunoprecipitation efficiency across samples, we computed a loading normalization factor *L_s_* for each sample *s*, defined as the median log2 PPM of all identified Mediator subunits *M* in that sample:

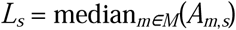

The normalized abundance of each protein was then obtained by subtracting this Mediator-derived normalization factor:

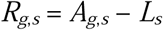

Finally, the bait-preferential score for each protein was defined as the difference between its normalized abundance in the MED16 IP and the mean of its normalized abundances in the other three Mediator bait IPs (MED1, MED12, CDK8):

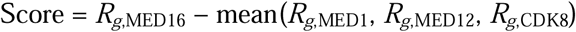

A positive score indicates enrichment in the MED16 IP relative to the other three Mediator baits. Candidate proteins of interest were further filtered by the following criteria: detection in at least two of three replicates of the target bait; maximum unique peptide count ≥ 2; detection in at least one additional Mediator bait; exclusion of Mediator/Pol II subunits, ribosomal proteins, histones, keratins, and common contaminants; and annotation with transcription or DNA-binding functions using GO terms from org.Hs.eg.db and GO.db, requiring either transcription regulator activity or DNA binding combined with a transcription-related biological process.

### Generation of *Med16* KO cell line using CRISPR-Cas9

To generate *Med16* knockout cell lines, the pX330-mCherry plasmid, containing the CRISPR-Cas9 machinery and target-specific guide RNAs, was used. A guide RNA sequence as reported in a previous study(77) was cloned into the pX330-mCherry vector. Cells were transfected with the pX330-mCherry plasmid using Lipofectamine 2000 (Invitrogen) according to the manufacturer’s protocol. After 48 hours, single mCherry-positive cells were sorted by fluorescence-activated cell sorting (FACS) into individual well of 96-wells plate. The sorted cells were then cultured and expanded. Individual clones were screened for gene knockout by sequencing and Western blotting to confirm successful editing.

### Retrovirus infection

Stable cell lines with retroviral-mediated knockdown or overexpression were generated following the manufacturer’s protocol (Clontech) as described previously (13). Specific siRNA target sequences were designed using Thermo Scientific’s siRNA selection tools. Retroviruses were produced by co-transfecting 293T cells with pSiren-RetroQ (for knockdown) or pMSCV-puro (for overexpression) vectors along with the pCL10A1 helper plasmid, using Lipofectamine 2000 (Invitrogen). After 48 hours, the culture medium containing retroviruses was collected, supplemented with 20 μg/ml polybrene (Sigma-Aldrich-Aldrich), and filtered through a 0.45 μm filter. The filtered retroviruses were then added to target cells, which were subjected to spin infection at 2,500 rpm for 1.5 hours at 300. Twenty-four hours after infection, cells were selected with puromycin in 293T at 1 μg/ml and HeLa cells at 2.5 μg/ml.

### Luciferase reporter assay

293T cells or HeLa cells were plated in a well of 12-wells plate (90% confluency) and transfected with 150 ng of luciferase reporter plasmid along with 50 ng pRL-TK (Promega) using Lipofectamine 2000 (Invitrogen). At 36-48h post transfection, the cells were harvested and subjected to dual-luciferase reporter assays according to the manufacturer’s protocol (Promega)

### RNA-seq and data analysis

Total RNA was extracted from cells lysed with TRIZOL. We used a commercial RNA library kit to prepare the RNA-seq library according to the manufacturer’s specifications (VAHTS® Universal V8 RNA-seq LibraryPrep Kit) from Illumina. Briefly, mRNA was captured with mRNA capture beads before fragmentation. The RNA fragments were reverse transcribed and subjected to library amplification for Illumina (Novaseq 6000) sequencing.

For RNA-seq data analysis, the filtered clean reads were mapped to the human reference genome hg38 with HISAT2. Read counts for each gene were then quantified using HTSeq-count. Differential expression analysis was performed in R using DESeq2, which normalized the read counts and identified significantly differentially expressed genes.

### Motif analysis

Motif enrichment analysis was performed using the closest known homolog of UBP1: the Tcfcp2l1 (GSE11431) position weight matrix (PWM) from the HOMER motif database (v4.9.1). Sequences and the Tcfcp2l1 PWM were submitted to the FIMO algorithm (MEME Suite v5.5.8) to scan for enriched motifs. This analysis generated base pair-resolution motif enrichment scores and associated p-values across all queried sequences.

### In vitro transcription assays with immobilized template

The immobilized template assay was carried out based on the previous study(78). Biotin-labeled PCR products were generated from the HIV-1 reporter plasmid with biotin attached to the upstream end. These templates contain the HIV-1 core promoter region (−78 to +60) followed by the luciferase CDS. 6 μg of each biotinylated template was bound to 300 μg of Dynabeads M-280 Streptavidin (Dynal) according to the manufacturer’s instructions using a DynalMag™-2 (Dynal) for assistance. The immobilized templates were then resuspended in 100 μL of blocking buffer (20 mM HEPES, pH 7.9, 0.1 M KCl, 10% glycerol, 6 mM MgCl2, 2.5 mM DTT, 50 mg/mL BSA). After a 15-minute incubation at room temperature, 100 μL of nuclear extract from either WT, MED16-overexpressing, or *Med16* KO HT cells was added and incubated at 30°C for 30 minutes to allow formation of pre-initiation complex (PIC). The immobilized templates were washed three times with buffer D containing 0.1% Triton X-100. The proteins bound to the templates were eluted with SDS gel loading buffer and analyzed by 10% SDS-PAGE.

For in vitro transcription, an NTP mix (25 mM each) was added to stimulate transcription after PIC formation, and the reaction was incubated at 30°C for 1 hour. The supernatant, containing the transcribed RNA products, was collected for qPCR quantification. The immobilized templates were again washed three times with buffer D plus 0.1% Triton X-100, and the proteins bound to the templates were eluted with SDS gel loading buffer and subjected to 10% SDS-PAGE.

### Quantification and statistical analysis

For qPCR and luciferase assays, the means and SEMs were calculated from at least three independent experiments. Statistical significance was assessed using GraphPad Prism software, with data presented in graphs as means ± SDs. Student’s t-test was used for comparisons between two groups, with significance levels defined as \**p* < 0.05 and \*\**p* < 0.01.

---

**Figure S1.**
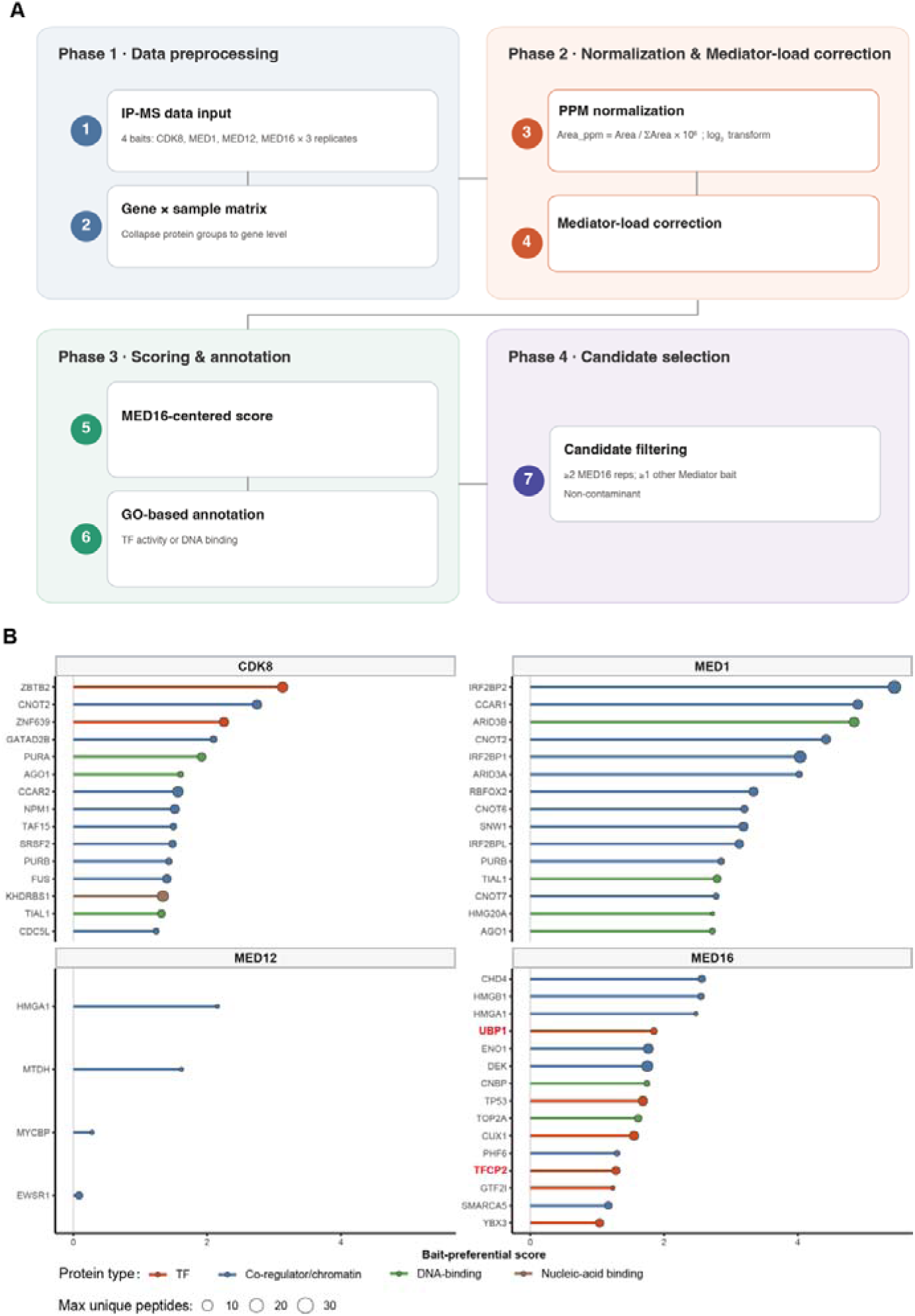
Identification of bait-preferential interacting proteins by Mediator subunit IP-MS analysis. (A) Workflow for identifying proteins preferentially associated with each Mediator bait (MED16 shown as an example). For each bait, proteins were normalized and corrected. A bait-preferential score was calculated for each protein. Candidate proteins were filtered by detection reproducibility, unique peptide count, and functional annotation. (B) Proteins identified as preferentially associated with each bait. For each bait, the top-ranked candidates are shown, with UBP1 and TFCP2 highlighted among the MED16-preferential candidates.

**Figure S2.**
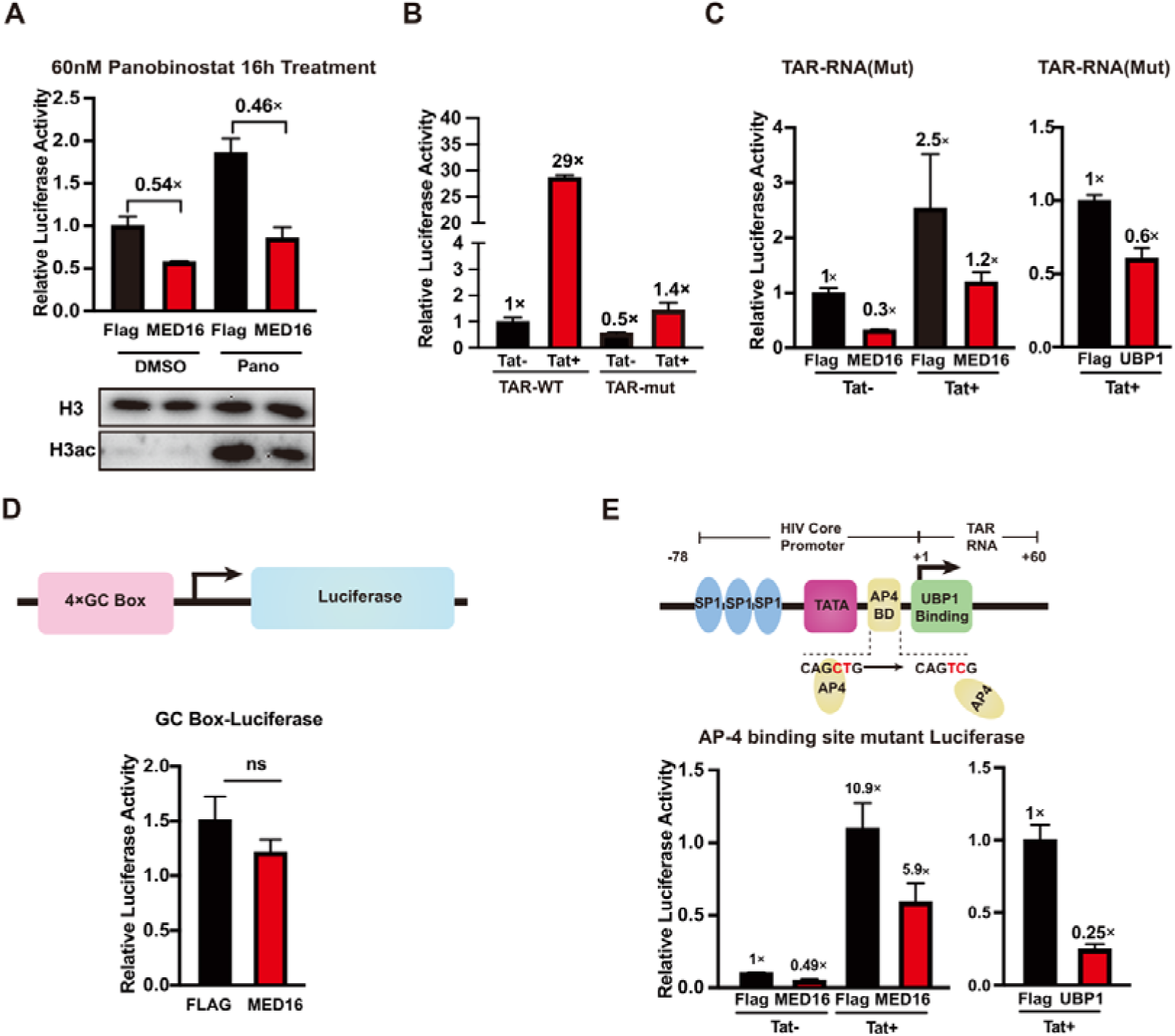
Identifying the MED16-controled regulatory elements at the HIV-1 promoter. 293T cells were transfected with 300 ng of either Flag control, MED16 overexpression, or UBP1 overexpression plasmid, along with the indicated reporter plasmids. (A) Cells were co-transfected with an HIV-1 reporter plasmid, then treated with or without 60 nM Panobinostat for 16 hours. Deacetylation inhibition was validated by western blot analysis (shown below). (B) Co-transfection with a TAR RNA mutant HIV-1 reporter and 50 ng of Tat overexpression plasmid, to assess the mutation’s effects. (C) Co-transfection with a TAR RNA mutant HIV-1 reporter to evaluate regulation by Flag control, MED16, or UBP1 overexpression. (D) Co-transfection with a 4×GC box reporter plasmid to examine MED16’s impact on GC box activity. (E) Co-transfection with an AP-4 binding site mutant HIV-1 reporter to assess MED16’s effect on the AP-4 site. The data are presented as the means ± SDs from n = 3 independent experiments; “ns”: no significant.

